# The overlooked role of muscle regeneration failure in post-implantation complications: a thorough investigation into mechanisms of recurrent urethral stricture

**DOI:** 10.64898/2026.07.16.738898

**Authors:** Alexey Fayzullin, Natalia Chepelova, Natalia Serejnikova, Nafisa Fayzullina, Mansur Mustafin, Andrei Bazarkin, Milena Bashkatova, Olga Drakina, Artem Antoshin, Yana Khristidis, Li Xue, Anyong Yu, Denis Butnaru, Evgeny Shpot, Eugene Bezrukov, Denis Chinenov, Peter Glybochko, Andrey Vinarov, Peter Timashev

**Author notes:** Dedicated to the memory of Professor Anatoly Shekhter.

## Abstract

Bioresorbable collagen membranes rarely achieve complete organ regeneration, often necessitating secondary operations. In this study, urethral defects were modeled in 60 Chinchilla rabbits; 30 were reconstructed using collagen membrane patches. Histological, immunohistochemical and in situ PCR analyses were performed at multiple time points up to 270 days post-implantation to assess inflammatory (TGF-β1, Wnt2, iNOS) and regenerative (collagen I/III, α-SMA, E-cadherin) markers. A biopsy from a patient with recurrent urethral stricture was analyzed using the same methodology. At three months after implantation, the mucosal layer had recovered, however, the underlying muscle layer remained incompletely regenerated. The muscle bundles were surrounded by α-SMA-positive myofibroblast-rich connective tissue with upregulated profibrotic markers. Comparable patterns of impaired muscle regeneration and high TGF-β1 expression were found in the human specimen. Our findings suggest that while muscle layer regeneration is essential for structural restoration, it may also trigger a sustained profibrotic cascade.

## INTRODUCTION

One of the major clinical challenges in reconstructive urology remains the treatment of extensive urethral strictures, particularly in patients with recurrent strictures. The current gold standard for managing such conditions is substitution urethroplasty using autologous tissue, most commonly buccal mucosa grafts. However, in long-segment urethral defects, the use of autologous grafts is associated with high recurrence rates and significant donor site morbidity, which can negatively impact patients’ quality of life and limit the feasibility of repeat procedures. ^1^

The design of novel biomaterial membranes is aimed to surpass regeneration potential of the autotransplantats in order to achieve restoration of long-segment urethral defects. Currently, both cell-based and acellular tissue-engineered constructs are being explored in clinical practice. However, cell-based technologies come with several limitations, including high costs and complex culturing procedures. In contrast, acellular matrices offer greater biocompatibility and broad application. Nevertheless, the indications for their use and the factors influencing successful reconstruction outcomes remain insufficiently understood, highlighting the need for further research and improvement of these materials.

Urethral reconstruction using decellularized matrices and tissue-engineered constructs has been actively investigated over the past decades. ^2^ These constructs are highly biocompatible but often exhibit low mechanical strength and limited reproducibility. In our previous studies, we demonstrated that decellularized matrices can support epithelial regeneration for up to three months. ^3–5^ However, the recurrence of stricture was observed during one-year follow-up. These findings underscore the need not only to develop mechanically robust and biocompatible implants, but also to assess their long-term performance in vivo.

Reconstruction of urethra remains one of the unsolved challenges of regenerative medicine. The regeneration factors are not only limited to epithelialization, vascularization and regulation of the inflammatory response but also include provision of an adequate biomechanical function. Disruption of these processes often leads to fibrosis, particularly beyond six months, as observed both in clinical practice and in experimental models. ^6–10^ Current evidence suggests that complications may be associated with the stimulation of epithelial-to-mesenchymal transition (EMT) in urethral epithelial cells by transforming growth factor beta 1 (TGF-β1), leading to their transformation into fibroblasts. This process results in excessive collagen synthesis and fibrosis, which narrows the urethral lumen. Moreover, decreased expression of E-cadherin, along with increased levels of vimentin and α-smooth muscle actin (α-SMA) - markers of mesenchymal cells, has been shown to correlate with stricture severity. ^11^ However, the EMT does not explain the specific long-term occurrence and predominantly central localization of the post-implantation stricture, which leaves the search for the primary cause of fibrosis open. Contributing factors under discussion include the effects of urine exposure, insufficient vascularization and excessive foreign body reaction. The limited understanding of the molecular mechanisms involved in inflammation and regeneration, continues to hinder the development of effective bioequivalents.

Based on our previous studies, we developed a mechanically robust collagen membrane that was initially evaluated in a subcutaneous model ^12^, later applied in dentistry in combination with lactoferrin ^13^ and successfully tested for tympanic membrane repair. ^14^ Comparative analysis with commercially available matrices demonstrated the advantages of the membrane in promoting tissue regeneration and modulating the inflammatory response. These findings provided the rationale for further investigation of the membrane in the context of urethral reconstruction.

The purpose of this study was to investigate the mechanisms of regeneration and fibrotic remodeling of the urethra during long-term follow-up period after implantation of the robust collagen membrane. Particular attention was given to the analysis of biointegration, epithelialization and connective tissue formation, with the goal of identifying molecular markers presenting targets for guided tissue regeneration.

## MATERIALS AND METHODS

### Fabrication and characterization of the collagen membrane

The procedure for fabricating collagen membranes has been described in the previous publication ^12^. Collagen was extracted from bovine Achilles tendons by homogenization in 0.5 M acetic acid followed by partial hydrolysis with 0.1% pepsin for 8 hours. The collagen was then precipitated using a 12% NaCl solution, redissolved and dialyzed against 0.5 M acetic acid for 72 hours. The final collagen concentration was adjusted to 5 mg/mL using acetic acid. Porous membranes were fabricated by electrophoretic deposition at 60 V for 20 minutes, followed by lyophilization. The membranes were treated with 99% isopropanol, dried, chemically cross-linked with a 0.03% glutaraldehyde solution, mechanically perforated using a mesoroller and lyophilized again at –40 °C.

The mechanical properties of the membranes were assessed using a Mach-1 v500csst testing system (Biomomentum Inc., Canada) after overnight incubation in PBS at 4 °C. Uniaxial tensile testing was performed on samples 30 × 5 mm at a strain rate of 0.1 mm/s until fracture. Ultimate tensile strength, elongation at rupture and Young’s modulus were determined.

The microstructure of the membranes was examined using scanning electron microscopy (SEM) with an EVO LS10 microscope (Zeiss, Germany) under low vacuum conditions (70 Pa, 21 kV, 40–70 pA). Sample preparation included rinsing, transverse sectioning with a microtome blade and air-drying.

### In vitro studies

To assess the indirect cytotoxicity of the collagen membranes in vitro, an Alamar Blue reduction assay was performed using human umbilical cord-derived MSCs. Membrane extracts (three fragments of 1 cm² per 1 mL of culture medium, incubated for 24 hours at 37 °C) were centrifuged, and the supernatants were added to MSC monolayers in 96-well plates. Serial dilutions of the extracts were tested in DMEM/F12 medium supplemented with 100 U/mL streptomycin, 100 μg/mL penicillin, 1% GlutaMAX (Gibco) and 5% fetal bovine serum (HyClone). Sodium dodecyl sulfate (SDS) solution was used as a positive control, while untreated culture medium served as a negative control. After 2 hours of incubation with Alamar Blue, fluorescence was measured using a Victor Nivo microplate reader (PerkinElmer, USA; excitation 580/20 nm, emission 625/30 nm). DNA content in the wells was quantified using the PicoGreen reagent after three freeze–thaw cycles (30 minutes each), followed by fluorescence measurement (excitation 480/30 nm, emission 530/30 nm).

Cell viability (3T3 fibroblasts and MSCs, seeded at 3 × 10⁴ cells/cm², cultured for 7 days) on collagen membranes (n = 5 per group) was visualized by Live/Dead staining (Calcein-AM 0.5 mg/mL, Sigma-Aldrich; propidium iodide 1.5 μM, ThermoFisher; Hoechst 33258 0.004 mg/mL, Thermo Scientific), followed by analysis with LSM 880 confocal microscope (Zeiss, Germany). Metabolic activity and DNA content were assessed as described above.

### Experimental model

All animal experiments were conducted in accordance with the international ARRIVE guidelines and approved by Local Ethics Committee of Sechenov University. The study was conducted on male Chinchilla rabbits (n = 60), aged 8–12 months and weighing 2.5–3.5 kg. All animals had fully developed genitourinary systems, without signs of congenital anomalies, urethral or perineal trauma, or infectious or inflammatory diseases. The animals were randomized into experimental (n = 30) and control (n = 30) groups. In the experimental group, urethral resection was performed followed by implantation of the collagen membrane. Animals in the control group underwent a sham operation without implantation. No food or water restrictions were applied prior to surgery. Sedation and surgical anesthesia were achieved using intramuscular medetomidine followed by intravenous propofol. Antibiotic prophylaxis (enrofloxacin), analgesia (ketoprofen) and local lidocaine infiltration were administered. All procedures were performed under sterile conditions. Animals were placed in the lithotomy position, and the surgical site was disinfected three times using a 1% iodopyrone solution and a chlorhexidine–alcohol mixture (1:1). A perineal incision was used to access the proximal penile and distal bulbar segments of the urethra. A 2.0 cm × 0.5 cm section of the dorsal urethra was excised. The resulting defect was repaired with a size-matched, individually shaped collagen membrane. The membrane was secured to the tunica albuginea of the corpora cavernosa with interrupted sutures (Vicryl 6-0), and its edges were anastomosed to the urethra with a continuous suture (Vicryl 6-0). After implantation, the surgical site was irrigated, hemostasis was achieved using a bipolar coagulator, and the wound was closed with interrupted sutures (Vicryl 5-0). A 1% iodopyrone solution was applied, followed by placement of a sterile dressing (Fig. S1). The urethra and catheter were secured to the anterior abdominal wall using a self-adhesive bandage. After recovery from anesthesia, the animals were placed in individual cages. Postoperative monitoring and wound healing assessment were carried out daily for the first 10 days, including wound cleaning and treatment with a 1% iodopyrone solution. Antibiotic therapy with enrofloxacin (2.5%, 0.2 mg/kg, intramuscularly) was administered for 7 days. Anti-inflammatory therapy was provided with subcutaneous ketoprofen (3 mg/kg). To ensure urinary drainage and prevent acute urinary retention, daily flushing of the urethral catheter with 0.9% NaCl solution was performed. The catheter was removed on day 7, after which all animals demonstrated complete restoration of spontaneous urination.

### Retrograde urethrocystography

Prior to euthanasia on post-operative days 15, 45, 90, 180 and 270, all animals underwent retrograde urethrocystography under pharmacological sedation with propofol (4–6 mg/kg). Up to 20 mL of a 38% urografin solution was retrogradely instilled through the urethral catheter while radiographic imaging was performed with a ¾ oblique projection to visualize the passage of contrast medium into the bladder. (Fig. S2)

### Histological evaluation of rabbit urethras

Animals were euthanized after the urethrocystography, six rabbits from each group (experimental and control) were sacrificed at the following time points: postoperative days 15, 45, 90, 180 and 270. Euthanasia was performed by administering Zoletil 100 (Vibrac, Carros, France; 60 mg/kg). The urethra and adjacent tissues from the reconstruction area were resected for histological evaluation.

The tissues were fixed in 10% neutral buffered formalin, underwent standard histological processing and were embedded into paraffin blocks. 4–5 µm thick sections were stained with hematoxylin and eosin, Mallory’s trichrome and Picrosirius red to visualize collagen fibers (Fig. S3-S4).

For immunohistochemical analysis, formalin-fixed paraffin-embedded tissue sections were deparaffinized and incubated with 3% hydrogen peroxide for 10 minutes to block endogenous peroxidase activity. A blocking solution (Cell Marque, USA) was used to prevent nonspecific antibody binding. Staining was performed using mouse monoclonal primary antibodies against α-smooth muscle actin (α-SMA; A2547, Merck, USA), inducible nitric oxide synthase (iNOS; MA5-17139, Thermo Fisher, USA) and E-cadherin (13-1700, Thermo Fisher, USA). Visualization was carried out using goat-derived secondary antibodies conjugated with horseradish peroxidase (HRP; G-21040, Invitrogen, USA), followed by chromogenic detection with diaminobenzidine (DAB) and counterstaining with hematoxylin.

Histological samples were digitized using a Hamamatsu NanoZoomer S20 slide scanner (Hamamatsu, Japan) at 400× magnification. Digital slide images were analyzed using the NDP.view 2 software (Hamamatsu, Japan). Polarized light microscopy of Picrosirius red-stained sections was performed using a LEICA DM4000 B universal light microscope equipped with a LEICA DFC7000 T digital video camera.

### In situ PCR

Formalin-fixed paraffin-embedded sections were deparaffinized and underwent antigen retrieval at pH 9.0 and 70 °C for 40 minutes. Slides were then incubated in 96% ethanol for 10 minutes and air-dried. The PCR reaction mixture with specific primers (Table 1), Taq polymerase, buffer and a fluorescently labeled nucleotide (sulfo-Cyanine5.5-dUTP; Lumiprobe, 2071-50nmol, USA), was applied directly onto the tissue sections and covered with a coverslip. Amplification was carried out using a flatbed thermocycler under the following conditions: denaturation at 95 °C for 15 seconds, annealing at 60 °C for 15 seconds and extension at 72 °C for 30 seconds, for a total of 30 cycles over 1 hour.

**Table 1.**
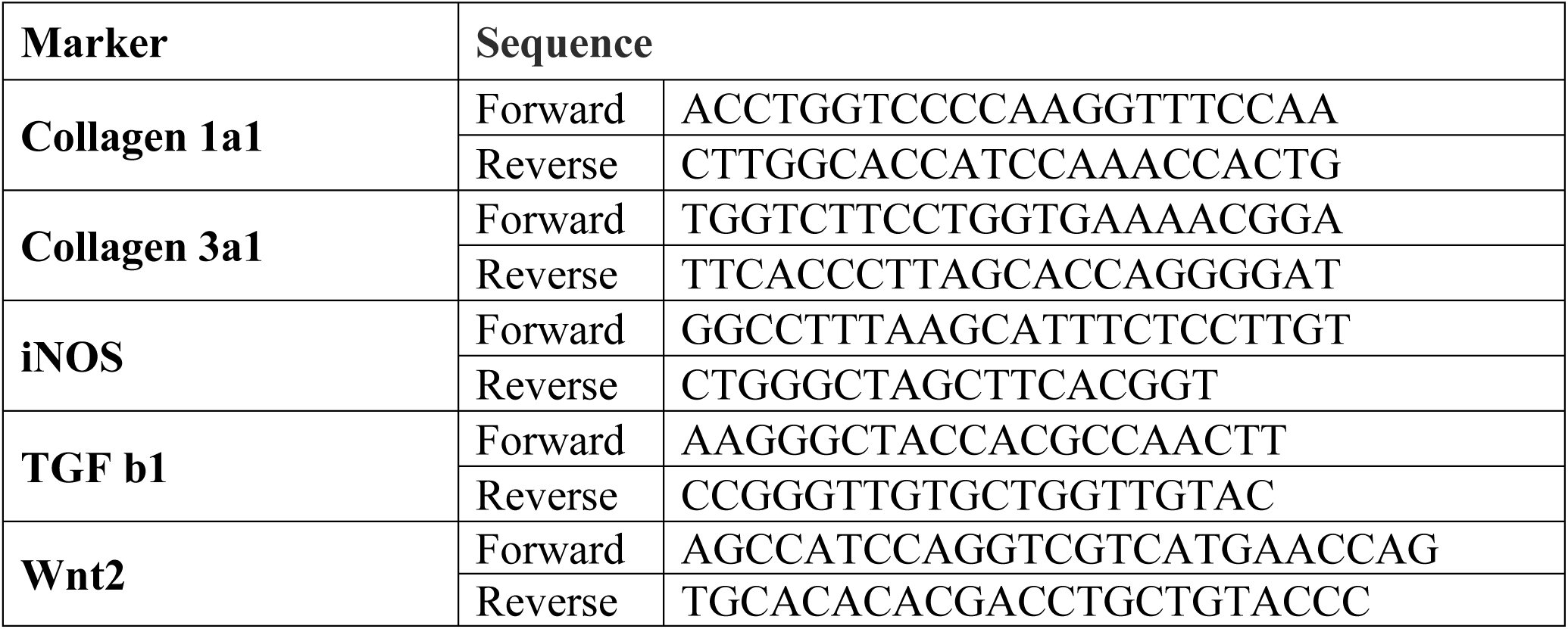
Primers used for in situ PCR in rabbit tissue.

Upon completion of the PCR reaction, the coverslips were removed using acetone, and the slides were rinsed in 96% ethanol for 10 minutes. Nuclear counterstaining was performed using DAPI. The samples were then analyzed using an Olympus FV3000 confocal microscope (Olympus, Tokyo, Japan).

### Morphometry

In histological sections stained with H&E and Mallory’s trichrome, a semi-quantitative scoring system was used to assess the severity of the inflammatory response (macrophage, lymphocyte, neutrophil and eosinophil infiltration, as well as fibrotic replacement of the collagen membrane) and parameters of tissue regeneration (epithelial restoration, membrane degradation and resorption, cellular infiltration into the implant structure, vascular network formation and overall tissue recovery). Evaluation was performed using a 5-point semi-quantitative scale: 0 — absence of the feature, 4 — strong manifestation (Table S1, Table S2, Table S3, Table S4, Table S5, Table S6, Table S7, Table S8, Fig. S5).

Quantitative analysis of the markers of collagen production (collagen types I and III) and inflammation (TGF-β1, WNT2, iNOS) was performed by calculating the portion of the positively stained cells within fields of view measuring 100 × 100 μm, with muscle bundles positioned at the center of each field (0 – no visible cells, 1 - < 25% of cells, 2 - >25% and <50% of cells, 3 - >50% and <75% of cells, 4 - >75% of cells). Evaluation was carried out in 10 fields of view per sample for each marker studied (Fig. 4).

### Morphological and molecular analysis of urethral tissue from a patient with recurrent stricture

To investigate the relevance of the identified mechanisms of urethral dysregeneration in clinical conditions, we analyzed urethral tissue samples obtained from a patient with recurrent stricture. A 50-year-old male patient presented with a recurrent stricture of the bulbar urethra. He previously underwent a two-stage buccal mucosa urethroplasty in 2016, followed by a repeat urethral reconstruction using buccal mucosa graft in 2025. Urethral tissue sampling was performed after obtaining written informed consent. The samples were stained with hematoxylin and eosin (H&E), Mallory’s trichrome and Picrosirius red (Fig. 5). Immunohistochemical (IHC) analysis was performed using antibodies against α-smooth muscle actin (α-SMA; A2547, Merck, USA) and in situ PCR (Table 2) was carried out according to the methodology described above.

**Table 2.**
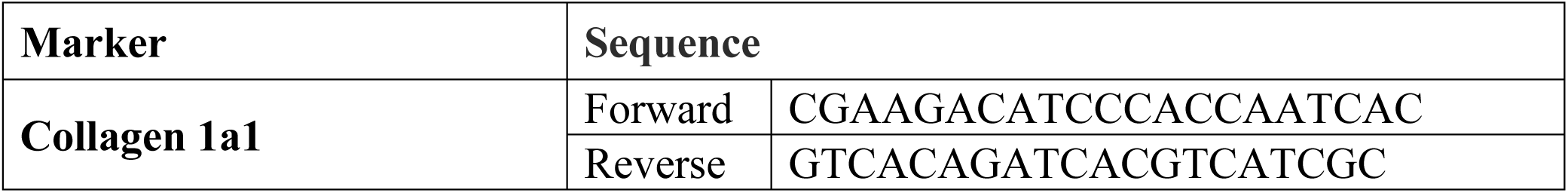
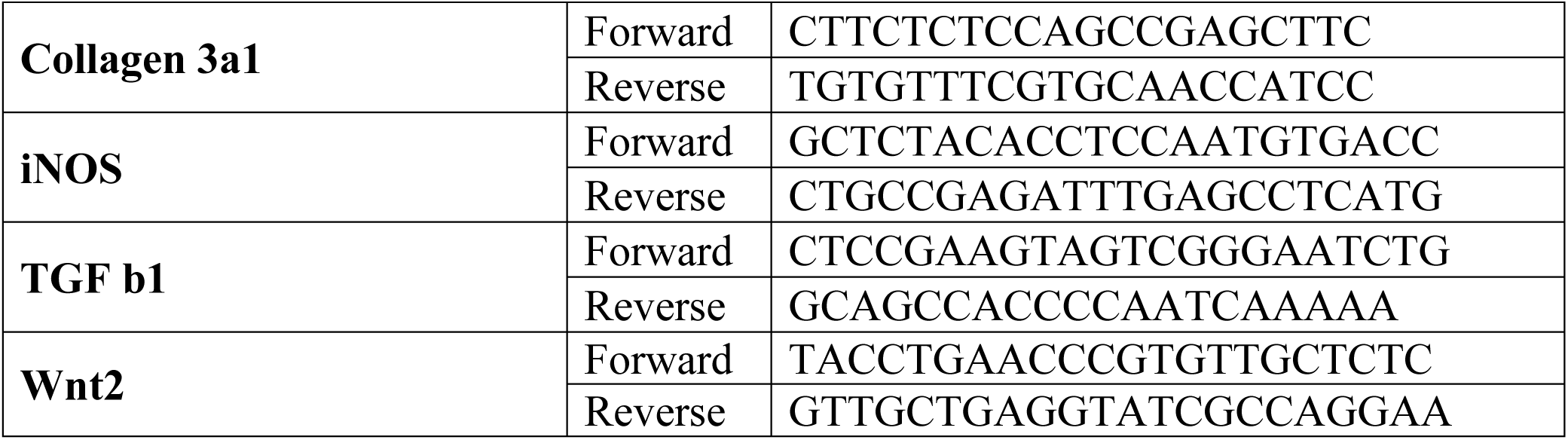
Primers used for in situ PCR in human tissue.

Tissue sections were digitized using the Hamamatsu NanoZoomer S20 histology slide scanner (Hamamatsu, Japan) at 400× magnification. Digital slide images were analyzed using NDP.view 2 software (Hamamatsu, Japan). Samples processed for in situ PCR were visualized using an Olympus FV3000 confocal microscope (Olympus, Tokyo, Japan).

### Statistical Analysis

Statistical analysis was conducted using GraphPad Prism 8.00 for Windows (GraphPad Software, USA). Differences in histological scores between groups were evaluated using the non-parametric Kruskal–Wallis test followed by Dunn’s multiple comparisons test. Differences in quantitative marker expression levels between groups were analyzed using one-way ANOVA followed by Tukey’s post hoc test. p values ≤ 0.05 were considered statistically significant. Results were presented as column graphs showing median values (for histological scores) and mean ± SD (for marker expression). (Fig. 4, Fig. S5)

## RESULTS

### Characterization of the collagen membrane

The fabricated collagen membranes were dense and exhibited following mechanical properties: the Young’s modulus was 2.2 ± 0.5 MPa and the elongation at break reached 51 ± 12%. One side of the membrane was smooth, providing barrier function and high tensile strength, while the opposite surface contained small open pores designed to enhance cellular integration. Cross-sectional imaging revealed a fibrous, tightly packed collagen structure with predominantly unidirectional fiber orientation, contributing to the implant’s mechanical robustness (Fig. 1).

**Fig. 1.**
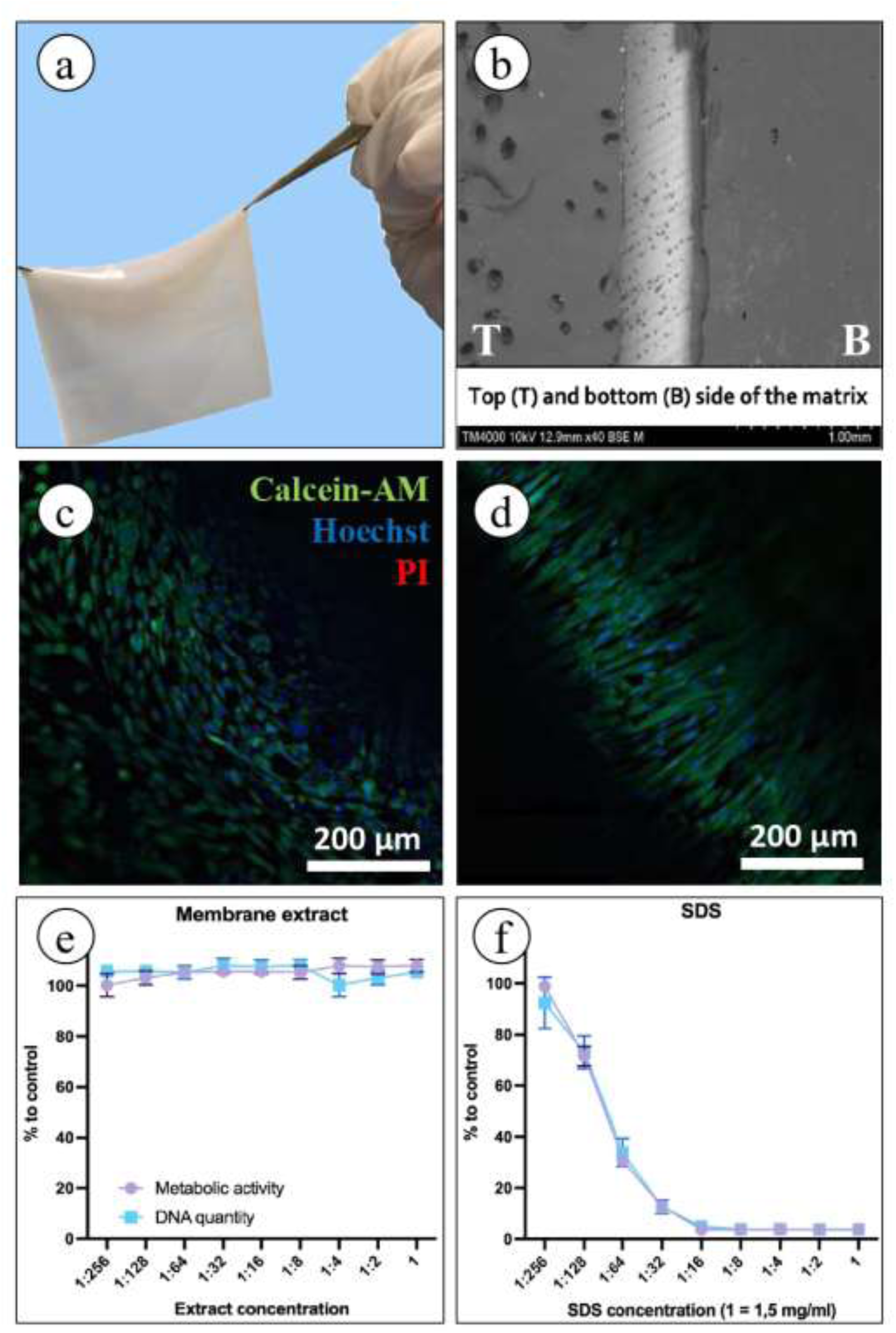
Characterization of the collagen membrane. (**A**) Fabricated collagen membrane. (**B**) Scanning electron microscopy (SEM) image of the membrane structure, the upper side of the membrane was perforated and the lower side was dense. (**C, D**) Direct cytotoxicity evaluation using Live/Dead staining of mesenchymal stromal cells (MSCs) and 3T3 fibroblasts (confocal microscopy): viable cells (green, Calcein-AM), dead cells (red, propidium iodide) and nuclei (blue, Hoechst). (**E, F**) Indirect cytotoxicity of collagen membranes toward MSCs, assessed via AlamarBlue (metabolic activity) and PicoGreen (DNA quantity) assays.

Extraction-based assays using AlamarBlue and PicoGreen demonstrated high biocompatibility of the membranes: cell viability and metabolic activity were not reduced by the application of chemical cross-linkers. Similarly, DNA content measured by the PicoGreen assay remained unchanged, confirming the absence of cytotoxic effects. In contrast, the positive control (sodium dodecyl sulfate - SDS) caused a marked decrease in cell viability. These findings indicate that the chemical cross-linking preserved the material’s functional characteristics without compromising its biocompatibility

Additionally, Live/Dead staining revealed effective cell adhesion and proliferation of both 3T3 fibroblasts and human umbilical cord-derived mesenchymal stromal cells on the surface of the collagen membranes. After six days of cultivation, all samples exhibited a high number of viable cells (green staining) with no evidence of dead cells (red staining). These results confirm that the developed collagen membranes support the adhesion and growth of various cell types, demonstrating excellent biocompatibility (Fig. 1).

### Retrograde urethrocystography

According to the retrograde urethrocystography, no signs of urethral stricture formation were observed in the experimental group during early follow-up periods (up to 90 days): the contrast medium filled the urethra evenly and flowed freely into the bladder, with no evidence of stenosis or extravasation. However, at later time points (180 and 270 days) progressive urethral narrowing accompanied by impaired urination was observed in some animals. On day 180, the contrast reached the bladder, but localized luminal narrowing was detected at the implantation site. By day 270, four animals exhibited pronounced stricture with contrast retention and marked reduction in urethral patency (Fig. 2).

**Fig. 2.**
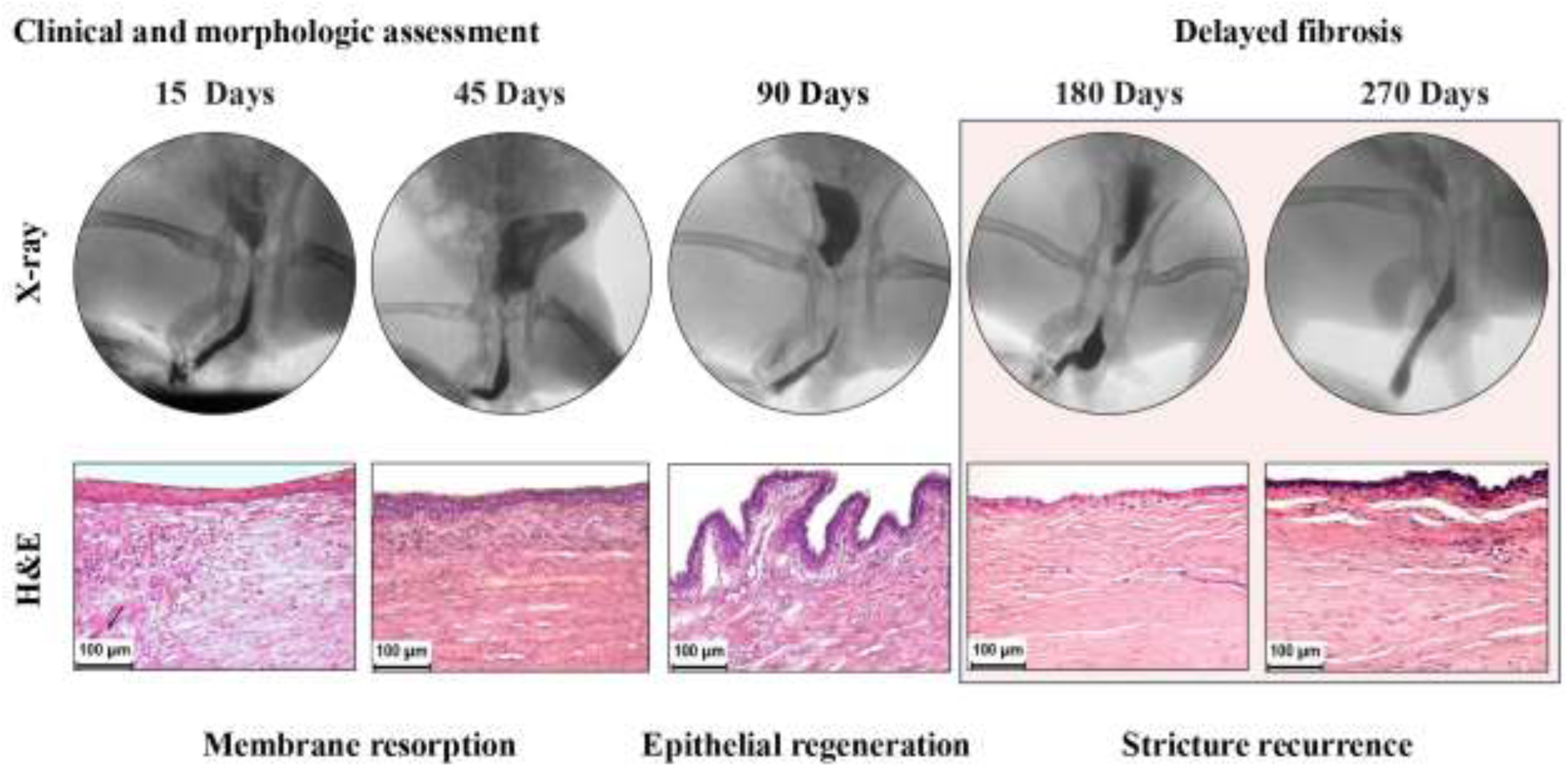
Clinical and morphological characterization of experimental outcomes at different time points. Up to day 45, no complications were observed. On day 90, no clinically significant abnormalities were detected on urethrocystography; however, early signs of tissue remodeling were identified in hematoxylin and eosin (H&E) stained slides. By days 180 and 270, urethral lumen narrowing and impaired contrast flow into the bladder were observed, accompanied by extensive fibrosis of the urethral mucosa at the implantation site, as confirmed by morphological analysis.

In the control group, signs of stricture formation appeared considerably earlier, by postoperative day 45. Four out of six animals showed signs of urinary obstruction, and retrograde urethrocystography demonstrated a typical radiographic pattern of urethral stricture formation.

### Histological analysis of the implantation sites in rabbits

On postoperative day 15, the experimental group demonstrated active epithelial regeneration, with the newly formed epithelium covering up to 50% of the implant surface. The regenerating epithelium exhibited features of low differentiation. In the control group, nearly complete epithelial coverage was also observed; however, areas of pronounced fibrosis and moderate lympho-macrophage infiltration were present. By day 45, the implant surface in the experimental group was fully epithelialized, although signs of functional differentiation remained limited. Immunohistochemical staining for α-SMA revealed the presence of myofibroblasts. In the control group, complete epithelialization persisted but was accompanied by pronounced fibrotic remodeling and a poorly differentiated epithelial layer.

By day 90, complete epithelialization was achieved in the experimental group, and no remnants of the collagen implant were detected. The replacement area showed mature epithelium without signs of inflammation. Mallory’s staining revealed clear evidence of implant resorption mediated by macrophages and accompanied by replacement with the host’s own connective tissue.

Immunohistochemical analysis of urethral sections was performed using antibodies against α-SMA, E-cadherin and iNOS to assess markers of fibrotic remodeling, epithelial integrity and inflammatory activity. α-SMA staining revealed smooth muscle bundles with a disorganized, chaotic arrangement in the central zone of the implantation site, indicating impaired regeneration of the urethral muscle layer (Fig 3A).

**Fig. 3.**
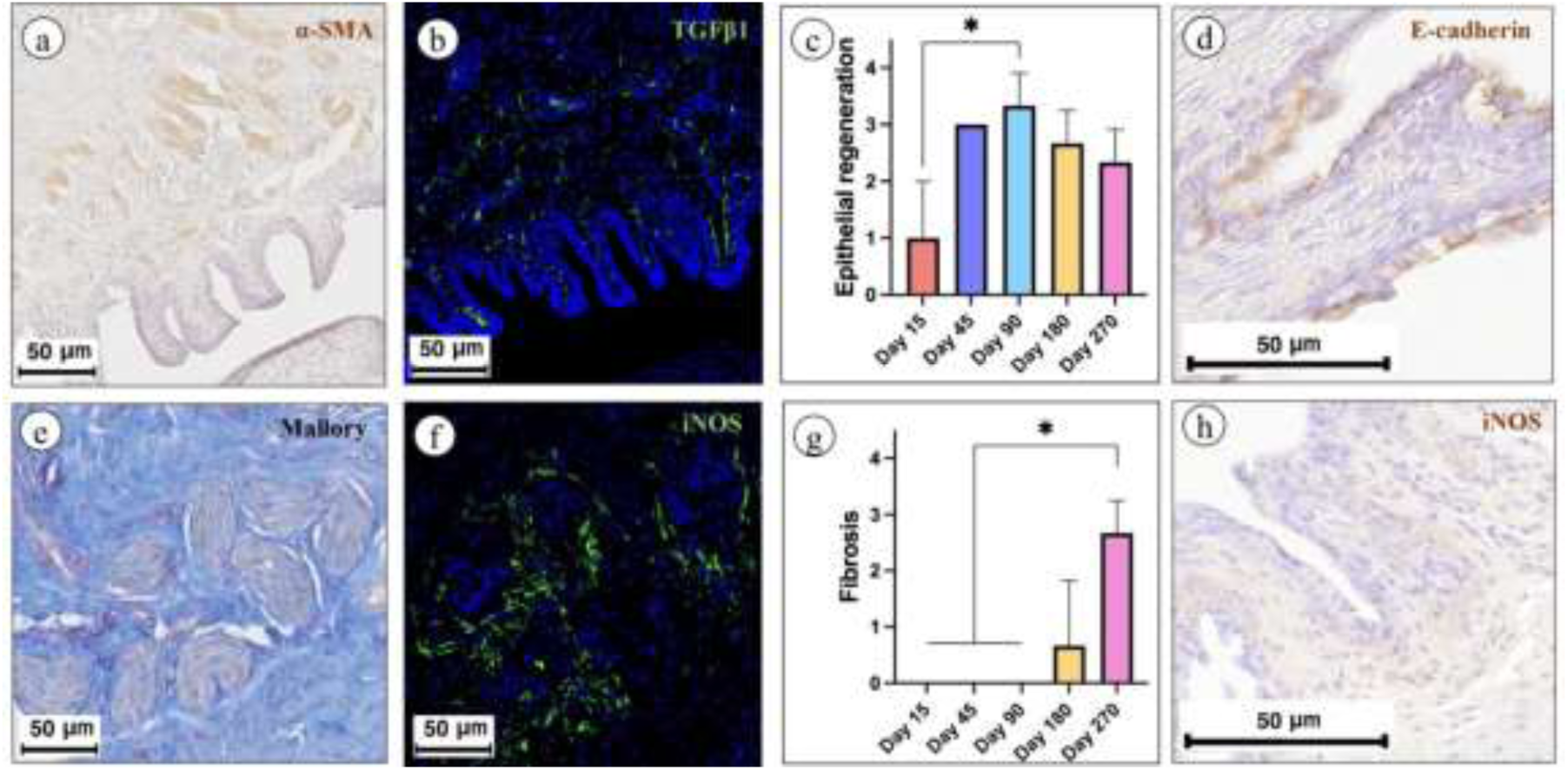
Morphological and molecular characterization of fibrotic remodeling of rabbit urethral tissue on day 90. (**A**) α-SMA-positive disorganized muscle fibers adjacent to the epithelial layer. (**B**) TGF-β1 expression in the perimuscular tissue environment. (**C**) Evidence of epithelial regeneration at all time points in the experimental group. (**D**) E-cadherin is a key marker of epithelial differentiation and intercellular adhesion, indicating a fully regenerated epithelium without signs of inflammation at 90 days after scaffold implantation. (**E**) Disorganized muscle fibers stained with Mallory’s trichrome. (**F**) iNOS expression surrounding muscle fibers. (H) Histologically confirmed fibrosis at late time points following scaffold implantation. (**H**) The absence of iNOS expression in the subepithelial zone indicates the absence of an active inflammatory response or a low level of local inflammatory activation in this area.

Expression of iNOS, a marker of inflammatory activity, was not detected in areas adjacent to the epithelium, suggesting that urine exposure is unlikely to be the primary trigger of late-stage fibrosis. However, iNOS-positive signals were observed in regions populated by α-SMA–positive cells, a pattern typically associated with responses to inflammation, injury or degenerative processes. It is well established that iNOS expression is induced in myocytes and infiltrating macrophages under conditions of tissue stress, supporting the interpretation that this localization reflects active tissue remodeling and highlights the involvement of the muscular component in fibrosis development.

Particular attention should be given to the expression of E-cadherin, which was strong and evenly distributed within the regenerated epithelium, indicating the restoration of intercellular junctions and the formation of a functional epithelial barrier. The combination of high E-cadherin expression and the absence of iNOS in the superficial layers suggests that neither urine leakage nor septic inflammation is the primary driver of stricture formation (Fig. 3D, H).

In contrast, the control group exhibited the formation of a dense fibrotic scar covered by a thin layer of poorly differentiated epithelial cells, along with extensive development of fibrous connective tissue.

At later time points (180 and 270 days post-implantation), approximately 25% of the mucosal surface within the implantation area in the experimental group exhibited dystrophic changes. In these regions, the urethral wall was lined with a thin epithelial layer and supported by a delicate connective tissue base lacking characteristic papillary structures. The underlying layers consisted of dense fibrotic scar tissue without signs of inflammation, composed of tightly organized bundles of collagen fibers and fibroblasts. The fibrotic component at the implantation site represented mature collagenous tissue, as confirmed by both Mallory’s trichrome and Picrosirius red stainings under polarized light.

Mallory’s staining revealed dense connective tissue bundles intensely stained blue, characteristic of collagen fibers. Polarized light analysis further clarified the degree of collagen maturity and fiber orientation: strong birefringence with predominant yellow-red hues indicated well-organized, densely packed collagen type I.

The presence of α-SMA in the form of scattered fragments rather than organized muscle layer supports the hypothesis that fibrosis may result from impaired regeneration of the muscle layer. In the control group, pronounced scar tissue and progressive urethral stricture remained evident at these time points.

### Morphometry

Analysis of dynamic changes in the morphological characteristics of the urethral mucosa at the implantation site revealed consistent and statistically significant shifts across several key parameters. In the experimental group, gradual loosening and resorption of the implanted membrane were observed, along with a progressive increase in its replacement by host connective tissue, as reflected by rising histological scores at later time points (day 180 and day 270).

Cell infiltration and vascularization of the membrane peaked at day 90, followed by a gradual decline, indicating the completion of the active implant integration phase. Simultaneously, epithelial regeneration and the formation of mucosal papillae progressively increased, reaching their maximum between day 90 and day 180. The levels of epithelial and papillary regeneration were significantly higher than those observed at earlier stages (day 15 and day 45), indicating restoration of the structural organization of the mucosa.

Fibrotic changes were minimal up to day 90 but significantly increased by day 180 and were most pronounced at day 270, reflecting long-term remodeling processes at the implantation site (Fig. S5).

### Assessment of molecular mechanisms of fibrosis development

Since histological and immunohistochemical analyses indicated that fibrotic changes in the urethra primarily developed in the central region of the implantation site, consistent with both the literature and our previous findings in human tissue, factors such as epithelial barrier dysfunction and insufficient vascularization were considered less probable to be the primary cause of the restructure. Therefore, an exploratory in situ PCR study was conducted to detect cell populations involved in excessive production of collagen and chronic inflammation.

Evaluation of COL1A1 (collagen type I) gene expression revealed a marked increase in signal intensity by day 90, with further upregulation at day 180. These results indicate a shift toward predominant accumulation of fibrotic collagen type I in the implantation zone by day 90; at a time when the urethral lumen was still patent, and mucosal regeneration had been morphologically completed. This suggests that pathological fibrosis began prior to the onset of clinical manifestations and that the excessive deposition of mature connective tissue in the central region of the implantation site served as its key substrate.

Analysis of TGF-β1 expression revealed a significant increase in fibrotically transformed areas at days 90 and 180, particularly around α-SMA-positive smooth muscle bundles. This pattern supports the conclusion that fibrosis progressed via a TGF-β1 dependent pathway and developed in regions of incomplete regeneration of the urethral muscle layer. By day 180, the dysregenerated muscle layer appeared to be the most pro-fibrotic tissue structure, indicating its potential key role in sustaining the fibrotic response.

An interesting finding was the localization of iNOS expression within muscle bundles on day 90, which coincided with a phase of active inflammation and tissue remodeling. However, by day 180, iNOS expression had nearly disappeared. Given that iNOS is not only a marker of inflammatory activation but also a key mediator of differentiation processes, its transient presence may be linked to early-phase responses, whereas its downregulation likely reflects the transition to a chronic fibrotic phase (Fig. 4).

**Fig. 4.**
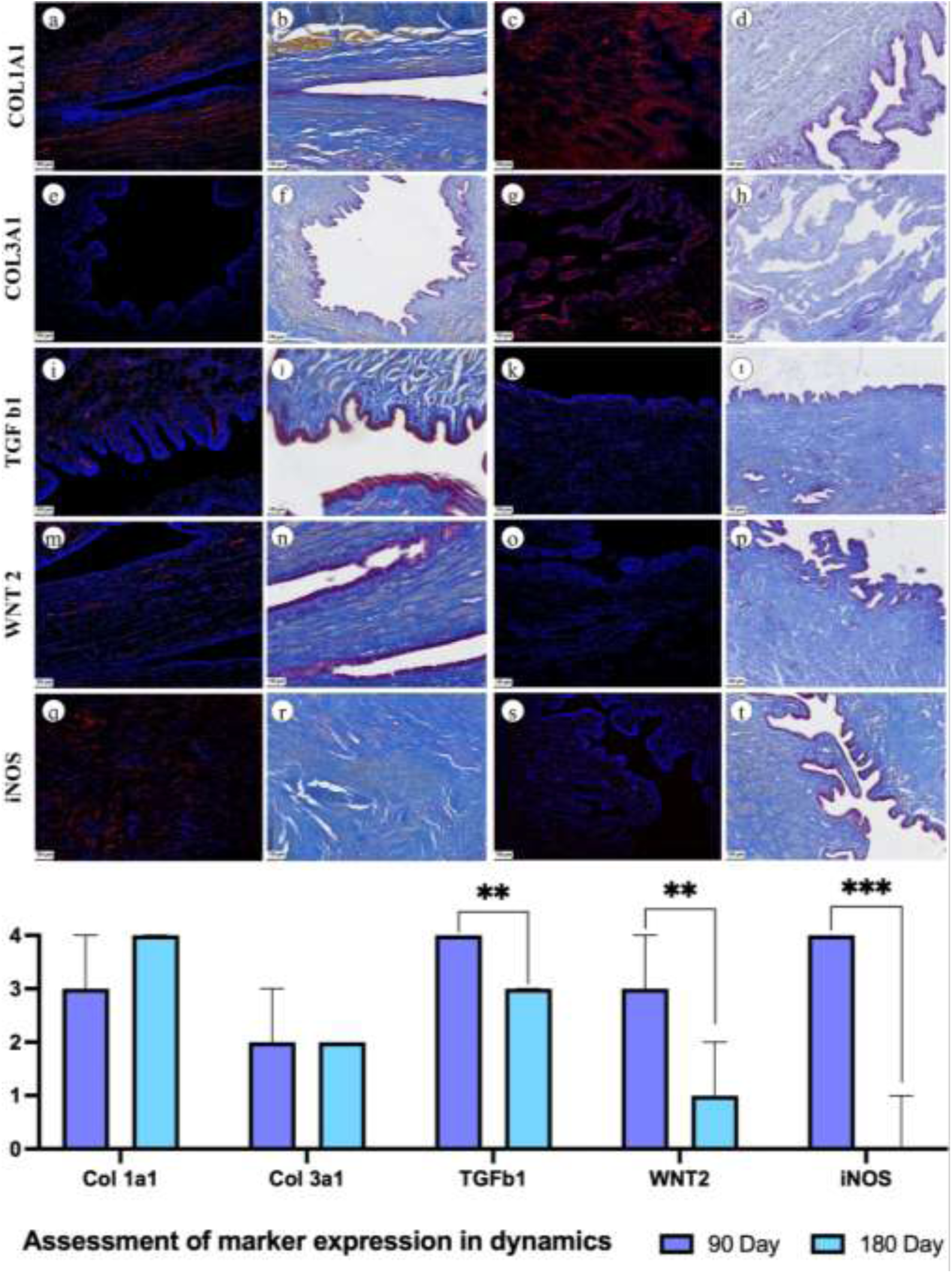
Comparison of in situ PCR results and Mallory’s staining at days 90 and 180 to analyze the expression dynamics of collagen types I and III (COL1A1 and COL3A1), WNT2, TGF-β1 and iNOS. Moderate decrease in the expression of COL1A1 at day 180 without statistical significance; collagen I remains an active component of the remodeling matrix even after 6 months, indicating that evidence of fibrotic activity persists. COL3A1 expression remains stable between 90 and 180 days. Collagen III continues to be involved in maintaining the structural integrity of the tissue, and the transition to mature collagen composition may be incomplete. High level of TGFβ1 expression on day 90 with clinically absent structure indicates the activity of the process, and its decrease on day 180 with preserved COL1A1 expression indicates the active process of fibrosis formation, which is clinically manifested by urethral stricture; Significant decrease in WNT2 expression by day 180, which indicates the weakening of signaling pathways of regeneration, which may be associated with the completion of reparative processes. High level of iNOS at day 90 indicates persisting inflammatory activity and may also be an early predictor of fibrosis formation. However, there is a sharp decrease in expression by day 180.

### Histological assessment of recurrent urethral stricture in a patient

Through comprehensive morphological, immunohistochemical and molecular analyses of urethral tissue in the experimental model, we identified a likely morphological agent of stricture formation – disorganized muscle bundles with strong α-SMA expression, accompanied by excessive TGF-β1 production in the surrounding stroma. To assess the clinical relevance of these findings and confirm the role of this mechanism in post-implantation stricture development, we performed a similar analysis of urethral tissue from a patient with recurrent bulbar stricture prior buccal mucosa urethroplasty.

Histological examination of the patient’s biopsy revealed that the ventral portion of the strictured segment was covered by stratified keratinizing epithelium of variable thickness. This morphology corresponded to the previously reconstructed neourethra formed from penile skin. Beneath the epithelium, a prominent layer of loose connective tissue was observed overlying the muscle layer. Immune cell infiltration was generally absent, appearing only in isolated areas beneath epithelial defects. Within the connective tissue, isolated α-SMA positive muscle bundles were identified, displaying weak iNOS expression. These structures exhibited irregular morphology, disorganized orientation and lacked clear architectural organization, consistent with a dysregenerative phenotype.

Mallory’s trichrome and Picrosirius red staining revealed dense collagenous matrix surrounding these bundles. Within the fibrotic tissue, spindle-shaped fibroblast-like cells were observed, exhibiting high TGF-β1 expression. These findings confirm the involvement of a TGF-β1-mediated signaling cascade in the formation of dense collagenous scar tissue and the subsequent development of urethral stricture (Fig. 5).

**Fig. 5.**
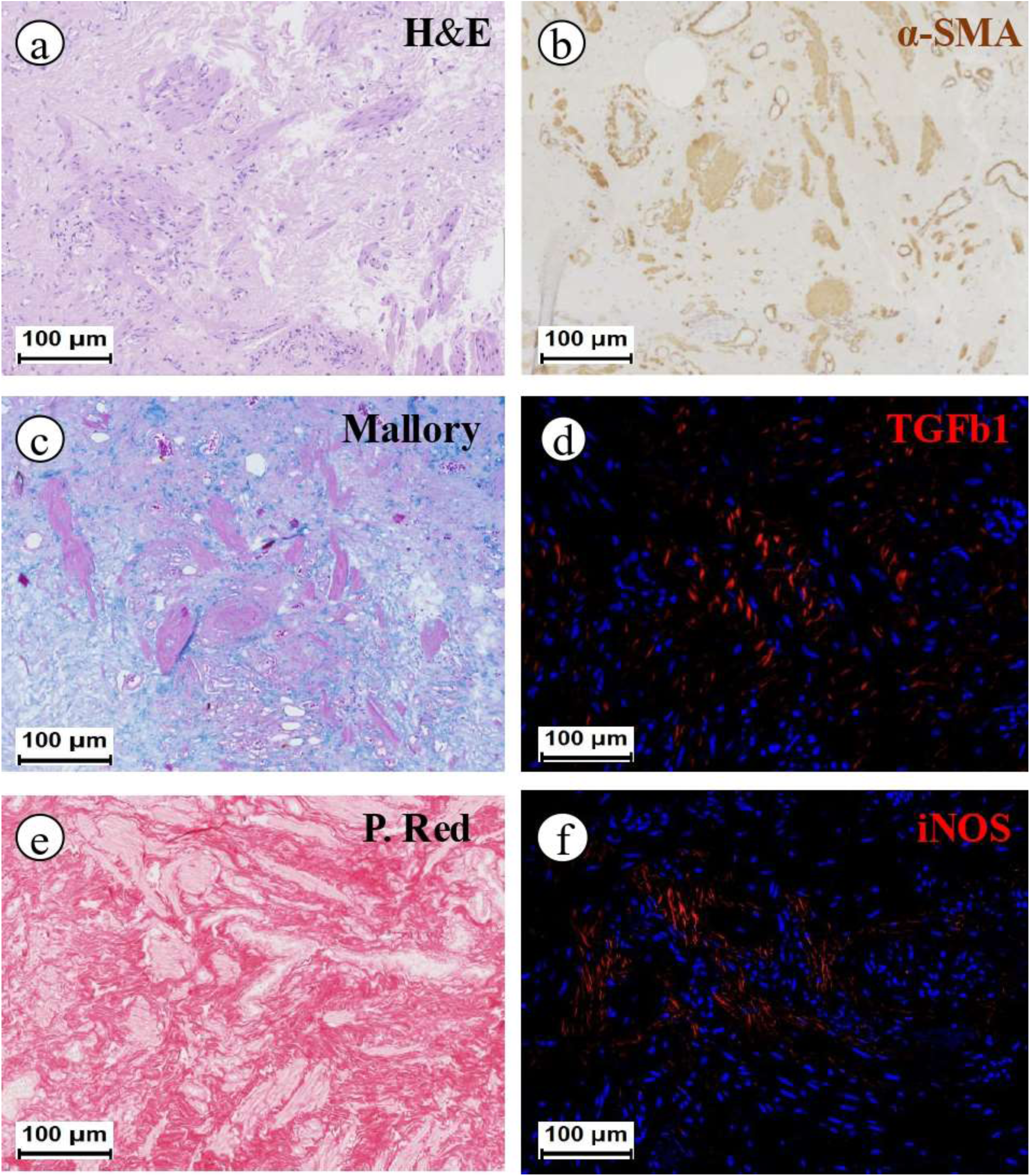
Histological and molecular characterization of a recurrent stricture. (**A**) Hematoxylin and eosin staining: cell nuclei - blue, cytoplasm and extracellular components – pink. (**B**) Immunohistochemical α-SMA staining: smooth muscle fibers and myofibroblasts were identified by brown staining. (**C**) Mallory staining: collagen fibers were represented by intense blue staining, muscle structures by pink staining. (**D**) Detection of TGF-β1 expression by in situ PCR - red signal in the zones of active fibrogenesis activity. (**E**) Picrosirius red staining: collagen fibers are detected as red, muscle fibers retain pale pink coloration. (**F**) Detection of iNOS expression by in situ PCR: positive signals are visualized as red in areas of inflammatory-destructive changes and myofibroblastic activity.

## DISCUSSION

In this study, we conducted an in-depth investigation into the mechanisms underlying post-implantation urethral stricture formation. Despite a substantial body of literature dedicated to reconstructive urology, several key questions remain unresolved: why do strictures emerge months after surgery rather than in the early postoperative period? Why does fibrosis originate at the center of the implantation site? Although some of these issues have been addressed in previous studies, including reports of delayed scar formation following urethroplasty ^6–10^, the underlying mechanisms – particularly those that could serve as potential therapeutic targets – remain poorly understood.

Within the framework of this project, we systematically ruled out mechanisms related to epithelial insufficiency and impaired vascularization. Morphological analysis demonstrated complete epithelial regeneration, supported by immunohistochemical expression of E-cadherin and the absence of epithelial defects. These findings align with previous studies that assessed epithelial barrier integrity using markers such as cytokeratins and laminin. ^15,16^ Furthermore, we observed no subepithelial immune cell infiltration, and morphometric analysis of blood vessels revealed no signs of vascular deficiency.

Two main hypotheses were considered: (1) a tissue reaction to membrane resorption, and (2) impaired regeneration due to localized abnormalities in cell migration or activity. The first hypothesis was deemed unlikely, as no significant numbers of macrophages or foreign body giant cells were observed in the implantation zone. The second hypothesis focused on fibroblast behavior, and in situ PCR analysis of collagen types I and III indicated local activation within the central zone without signs of peripheral migration from the tunica albuginea or defect edges.

The crucial finding was the colocalized expression of COL1A1 and TGF-β1 within connective tissue adjacent to disorganized α-SMA–positive muscle bundles. These regions, located centrally within the implantation site, represented zones of incomplete regeneration of the muscle layer – the only urethral wall component that had not recovered by day 90. Moderately elevated iNOS expression in these zones may reflect an attempt at cellular differentiation or a response to oxidative stress. ^17^ However, its disappearance by day 180 suggests exhaustion of protective mechanisms. Given their specific localization, expression profiles and morphological features, we conclude that these structures constitute the morphological substrate of post-implantation urethral stricture. Notably, similar features were also identified in clinical samples from a patient with recurrent stricture.

The potentially protective role of iNOS warrants particular attention: it is plausible that the downregulation of iNOS contributes to the acceleration of fibrotic processes. ^18^ This highlights the potential of nitric oxide donors as a promising strategy for the prevention and treatment of urethral strictures. Accordingly, the therapeutic relevance of nitric oxide donors in mitigating urethral fibrosis merits further investigation.

Taken together, these findings offer a novel perspective on the etiology of post-implantation complications. In designing implantable materials, it is essential to address not only epithelial regeneration but also guided regeneration of the muscle layer – resembling strategies employed in peripheral nerve tissue engineering. ^19^ In addition to bioengineering solutions, therapeutic approaches such as targeted modulation of TGF-β1 signaling – e.g., via agents like pirfenidone ^20^, potentially delivered locally – are emerging as relevant and promising avenues for future research.

Limitations of the present study include the use of a model of urethroplasty without preexisting fibrotic damage to the urethral tissue. As such, the findings require validation in models involving preformed scar tissue at the implantation site ^21–23^, as well as models employing tubularized implants. ^24–26^ In addition, the experiments were conducted on young animals, which is common in most comparable studies. ^27–29^ However, it is well established that tissue regenerative capacity is higher in young individuals than in aging organisms. Given the strong age-related association observed in urethral stricture disease ^30^, future studies using aged animal models are warranted to enable more accurate extrapolation of the results to clinical practice.

## CONCLUSIONS

In summary, we demonstrated that that incomplete regeneration of the urethral muscle layer following collagen membrane implantation is a key driver of late-onset complications and stricture formation. The central implantation zone, characterized by disorganized α-SMA-positive muscle bundles and sustained TGF-β1 expression, emerges as the primary morphological substrate of post-implantation complications. These findings were consistent across the rabbit model and human recurrent stricture tissue, underscoring their translational relevance. Our results suggest that muscle regeneration, while essential for structural repair, can paradoxically initiate and sustain a profibrotic cascade in the absence of proper tissue organization. Notably, this mechanism lies outside the prevailing theory of biological integration of implants, which typically emphasizes epithelialization, vascularization and inflammatory modulation as primary determinants of long-term success.

## Acknowledgments

The animal study was supported by the Russian Science Foundation (Grant No. 23-15-00481). The biomaterial design research was carried out within the framework of the state assignment of the Ministry of Health of the Russian Federation (№ NZAF-2024-0025). The researchers express their gratitude to Chopav Abdulaev and Yana Svetocheva for their valuable contributions to the modeling of urethral defects, performing implantations and providing attentive animal care during the follow-up period.

## Data availability

The data supporting the findings of this study are available within the paper and its Supplementary Information. The minimum dataset required to interpret, verify and extend the findings, including original histological images, morphometric measurements and source data for all graphs, is available from the corresponding author upon request.

## Author Contribution

Conceptualization, N.S., A.B., M.B., A.A., D.B., E.S., E.B., D.C., P.G., A.V. and P.T.; methodology, A.F., N.C., N.F., M.M., M.B., O.D., A.A., Y.K., L.X., A.Y. and E.B.; software, A.F., N.S. and O.D.; formal analysis, N.C., M.B., A.A. and D.C.; investigation, A.F., N.C., M.M., A.B., O.D., A.A., Y.K., D.B., E.S., D.C., P.G., A.V. and P.T.; resources, A.A., L.X., A.Y., A.V. and P.T.; data curation, O.D.; writing—original draft preparation, A.F., N.C., A.A., Y.K., E.S., D.C. and P.G.; writing—review and editing, N.S., N.F., M.M., A.B., M.B., O.D., L.X., A.Y., D.B., E.B., A.V. and P.T.; visualization, A.F.; supervision, D.B. and E.B.; project administration, E.S., P.G., A.V. and P.T.; funding acquisition, E.S., P.G., A.V. and P.T..

