## Supplementary files for "The overlooked role of muscle regeneration failure in post-implantation complications: a thorough investigation into mechanisms of recurrent urethral stricture"

**List of Supplementary Materials**

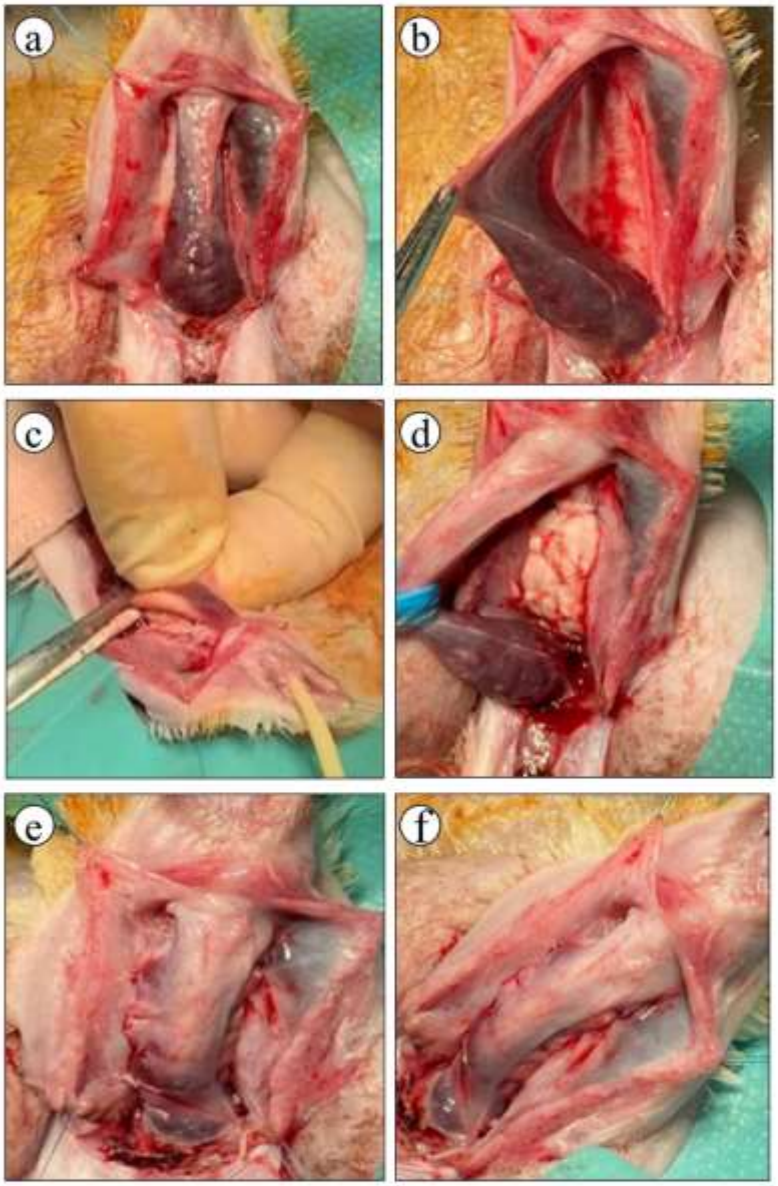

**Fig. S1. Surgical procedure. (A)** Surgical access to the proximal penile and distal bulbar urethra. (**B)** Mobilization of the distal bulbar urethra. **(C, D)** Fixation of the membrane to the ventral surface of the tunica albuginea of the corpora cavernosa and excision of the urethral segment**. (E, F)** Suturing the edges of the Janus-type membrane to the urethral margins at the defect site.

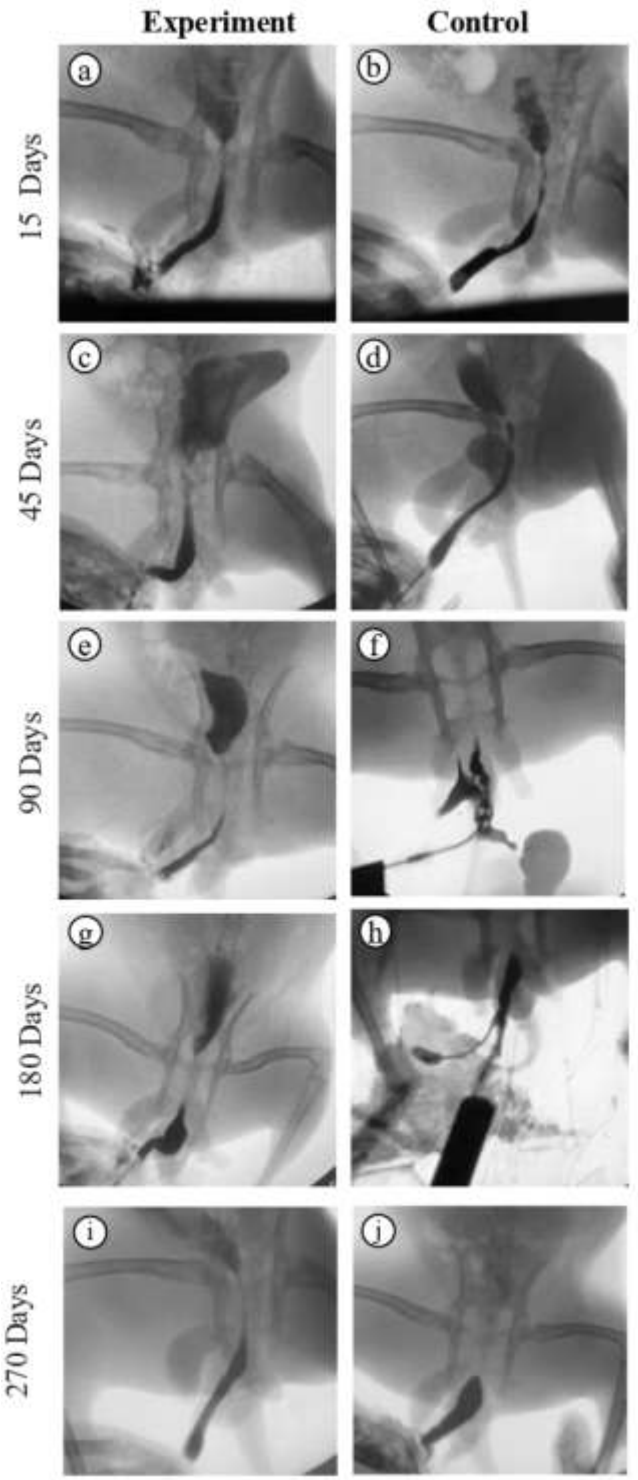


**Fig. S2. Retrograde urethrocystography: experimental and control groups at all time points. (A, C, E) In the experimental group, by 15-90 days after the**

**intervention, satisfactory urethral patency was observed: contrast agent freely passed through the urethra into the bladder, and the lumen was preserved. (G) However, starting from 180 days, clinically significant narrowing of the urethral lumen with impaired contrast flow into the bladder is visualized, which indicates the development of late urethral stricture. (I) On the 270th day a pronounced contrast passage disorder remains, confirming a persistent stricture change. (F) In the control group, a pronounced urethral patency disorder is detected already on the 90th day: the contrast agent is not visualized in the bladder, indicating the formation of a pronounced stricture. (H, J) This picture persists at 180 and 270 days of observation, which confirms persistent narrowing of the urethral lumen.**

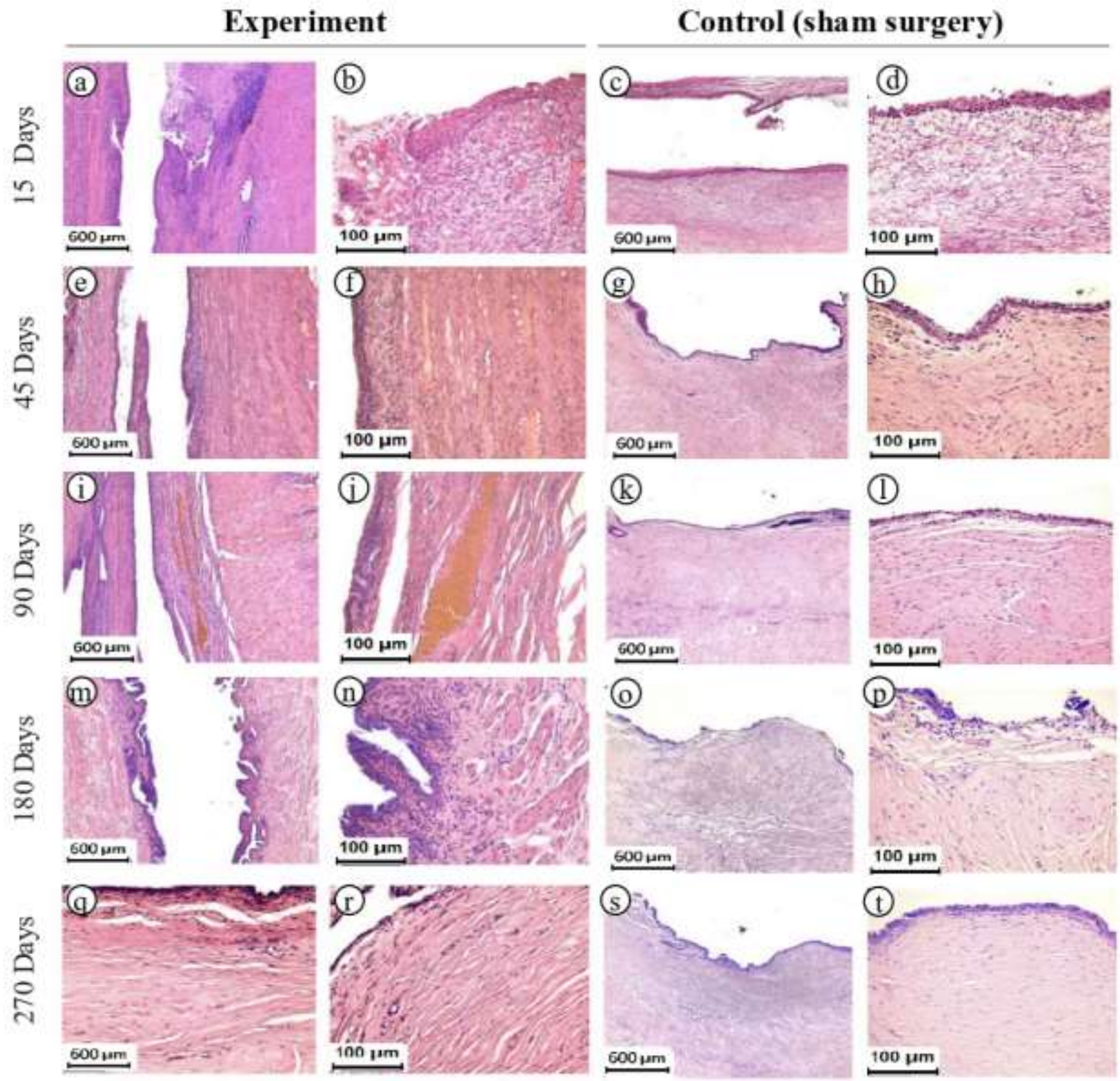


**Fig. S3. Hematoxylin and eosin staining: experimental and control groups at all time points. Experiment: (A, B) 15 days after surgery, partial epithelialization and initial resorption of the collagen implant with vessel ingrowth and cellular infiltration are noted. Microscopy reveals regeneration of the urothelium, active phagocytic reaction and formation of the transition zone between the implant and the urethral wall. (E, F) 45 days after surgery, the implant is**

**partially resorbed, its remnants are integrated into the tissue of a layered structure with fully formed urothelium. Areas with collagen fibers, smooth muscle bundles and newly formed vessels are noted. (I, J) 90 days after the operation, fully epithelialized fibrous tissue with large vessels replacing the implant is formed in the defect area. The surface is covered with mature urothelium with mucosal papillae forming. (M, N) 180 days after the operation, the defect area shows regenerated mucosa with the structure corresponding to the intact urethra, with urothelium, vessels and smooth muscle bundles. However, in some areas there are areas of pronounced fibrosis with thinning urothelium and predominance of fibrous scar tissue. (Q, R) 270 days after surgery in the area of the defect there is mucosa regeneration with urothelium of normal structure and moderate inflammatory infiltration. However, there is a pronounced fibrous thickening of the wall and a sharp narrowing of the urethral lumen due to the replacement of tissues with fibrous scar structure.**

**Control: (C, D**) After 15 days, the urothelium became multilayered and more mature, and a pronounced fibrosis developed in the connective tissue base of the mucosa. (**G, H**) After 45 days, the urethral mucosa in the defect area was completely epithelialized, but the urothelium remained poorly differentiated, heterogeneous in thickness and with pronounced vacuolar dystrophy. In the underlying connective tissue a pronounced fibrosis was formed with the presence of fibrotic scar tissue, lympho-macrophage infiltration and fibrotic vessels. (**K, L**) In 90 days the urethral mucosa in the defect area was represented by a thick layer of scar tissue, in some places with hyalinosis. The mucosal papillae were absent, and the scar was covered with a thin layer of dystrophically changed and in some places desquamated epithelial cells. (**O, T**) At 180 and 270 days, scars were detected in the defect area in all animals. The scars consisted of collagen fiber bundles located longitudinally, randomly or in the form of knots. The scars were covered with a thin layer of epithelium with dystrophic changes, there were also areas of epithelium desquamation. Lympho-macrophagal infiltration was mostly weak, but in some places near the scar and in the scar it was increased. In the mucous membrane at the border with the scar there were sometimes detected areas of edema.

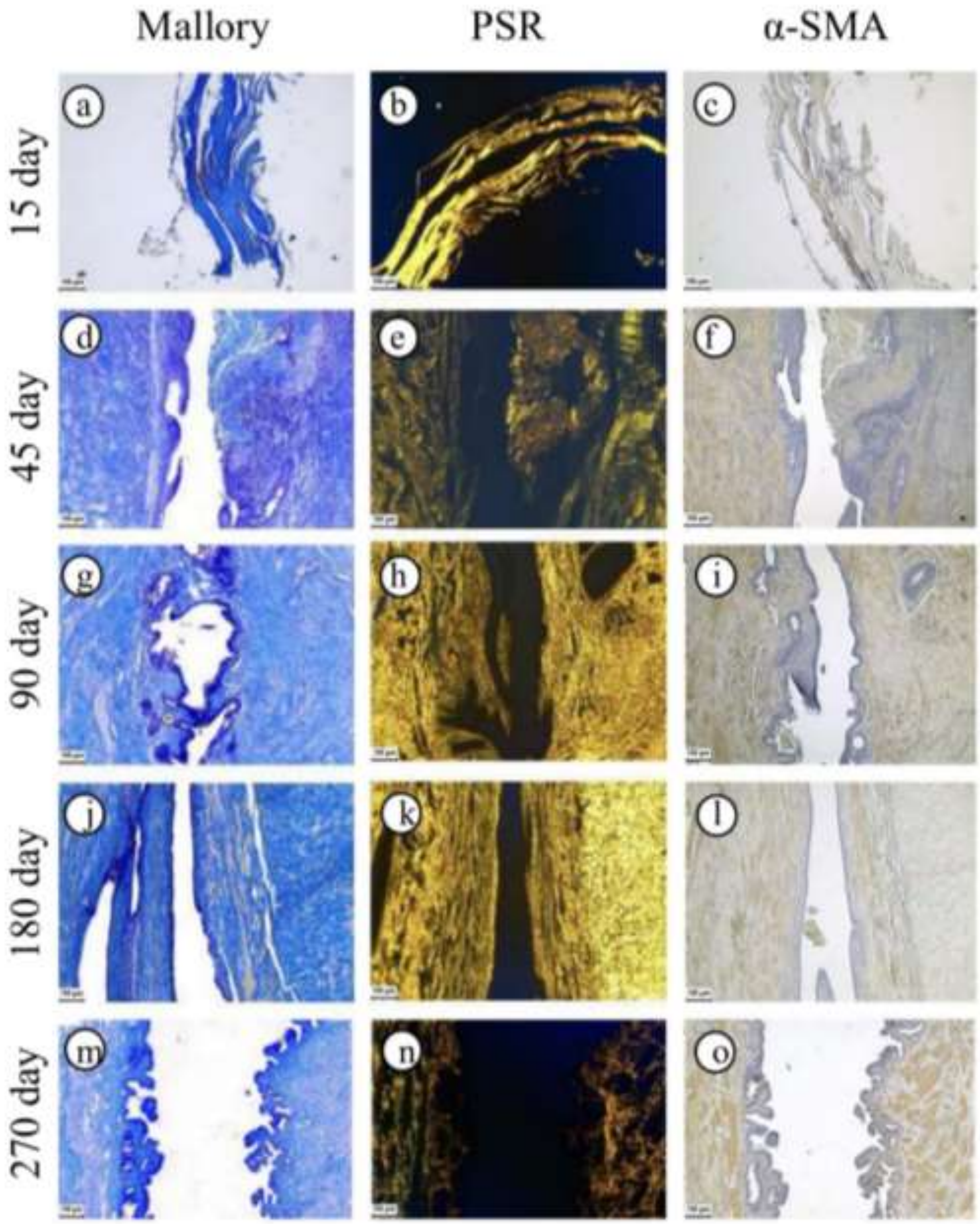


**Fig. S4. Mallory's trichrome, Picrosirius red and α-SMA staining: experimental group at all time points. Collagen fibers of intrinsic connective tissue are blue (Mallory), giving yellow anisotropy (Picrosirius red). Bundles of smooth muscle cells are colored pink (Mallory); α-SMA (α-smooth muscle actin) stain smooth muscle cells and myofibroblasts brown.**

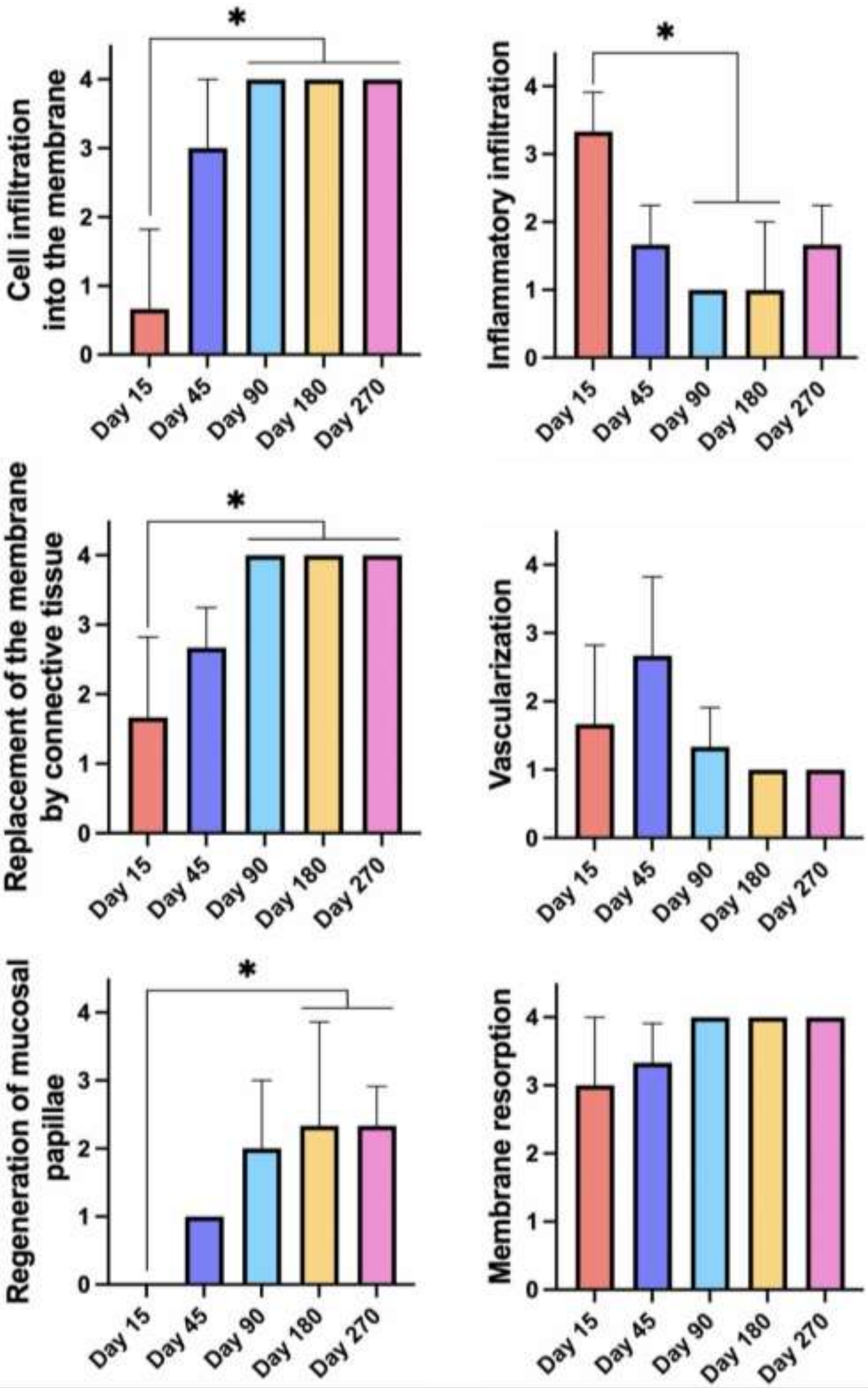


**Fig. S5. Morphometry: experimental group at all time points.**

**Table S1.** Scoring system for morphologic signs of **epithelial regeneration**

| | |
|---|---|
| 0 | non-dense single-layer epithelium |
| 1 | non-dense monolayer epithelium with several areas of multilayer epithelium |
| 2 | continuous single-layer epithelium with several areas of immature multilayer epithelium |
| 3 | split differentiated multilayered epithelium with areas of immature multilayered and unilayered epithelium |
| 4 | split multilayered differentiated epithelium |

**Table S2.** Scoring system for visual signs of **fibrosis**

| | |
|---|---|
| 0 | single bundles of collagen fibers, without formation of mature fibrous tissue |
| 1 | presence of immature fibrous tissue in subepithelial or submembranous zones |
| 2 | moderate amount of fibrous tissue with areas of organized collagen |
| 3 | pronounced fibrosis with predominance of mature collagen fibers |
| 4 | dense mature fibrous tissue replacing most of the area under study |

**Table S3.** Scoring system for morphologic signs of **cell infiltration into the membrane**

| | |
|---|---|
| 0 | cells are absent on the surface and inside the membrane |
| 1 | single cells on the membrane surface |
| 2 | small number of cells partially infiltrated into the membrane structure |

| | |
|---|---|
| 3 | moderate number of cells infiltrating the membrane over the whole area |
| 4 | massive ingrowth of cells into the entire membrane thickness |

**Table S4.** Scoring system for morphologic signs of **inflammatory infiltration at the implantation site.**

| | |
|---|---|
| 0 | absence of immune cell foci |
| 1 | single small foci of immune cells |
| 2 | few foci of immune cells |
| 3 | moderate diffuse infiltration with immune cells |
| 4 | significant diffuse infiltration with immune cells |

**Table S5.** Scoring system for morphologic signs **of replacement of the membrane by connective tissue**

| | |
|---|---|
| 0 | membrane is completely intact, without signs of replacement |
| 1 | initial signs of replacement in separate areas |
| 2 | replacement by connective tissue up to 50% of the membrane area |
| 3 | replacement prevails, membrane is preserved only fragmentarily |
| 4 | complete replacement of the membrane by connective tissue, membrane remnants are not visualized |

**Table S6.** Scoring system for morphologic signs of **vascularization**

| | |
|---|---|
| 0 | no vessels |

| | |
|---|---|
| 1 | single immature capillaries |
| 2 | moderate number of vessels, mainly immature vessels |
| 3 | numerous vessels, including mature vessels with a pronounced wall |
| 4 | abundant vascularization, mature vessels with perivascular structure predominate |

**Table S7.** Scoring system for morphologic signs of **regeneration of mucosal papillae**

| | |
|---|---|
| 0 | absence of papillae |
| 1 | single sparse papillae on the surface of the mucous membrane |
| 2 | papillae occupy 1/2 of the mucous membrane surface |
| 3 | papillae occupy 2/3 of the mucous membrane surface |
| 4 | papillae are formed on the whole surface of the mucous membrane |

**Table S8.** Scoring system for morphologic signs of **membrane resorption**

| | |
|---|---|
| 0 | membrane is completely preserved, no signs of degradation |
| 1 | initial signs of surface resorption |
| 2 | resorption affects up to 50% of membrane thickness or area |
| 3 | resorption is pronounced, membrane is thin or discontinuous |
| 4 | complete resorption, membrane is not detectable |
